# Salt-dependent conformational changes of intrinsically disordered proteins

**DOI:** 10.1101/2021.05.20.444991

**Authors:** Samuel Wohl, Matthew Jakubowski, Wenwei Zheng

**Affiliations:** Department of Physics, Arizona State University, Tempe, AZ 85287, USA; College of Integrative Sciences and Arts, Arizona State University, Mesa, AZ 85212, USA

## Abstract

The flexible structure of an intrinsically disordered protein (IDP) is known to be perturbed by salt concentrations, which can be understood by electrostatic screening on charged amino acids. However, an IDP usually contains more uncharged residues which are influenced by the salting-out effect. Here we have parameterized the salting-out effect into a coarse-grained model using a set of Förster resonance energy transfer data and verified with experimental salt-dependent liquid-liquid phase separation (LLPS) of 17 proteins. The new model can correctly capture the behavior of 6 more sequences, resulting in a total of 13 when varying salt concentrations. Together with a survey of more than 500 IDP sequences, we conclude that the salting-out effect, which was considered to be secondary to electrostatic screening, is important for IDP sequences with moderate charged residues at physiological salt concentrations. The presented scheme is generally applicable to other computational models for capturing salt-dependent IDP conformations.

**Graphical TOC Entry:** 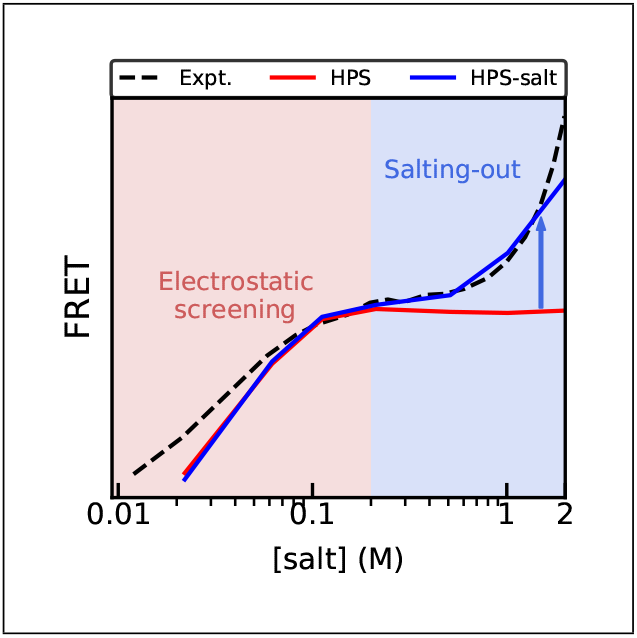

---

Emerging knowledge of intrinsically disordered proteins (IDPs) has shown us numerous examples of functions without a well-defined three-dimensional structure.^1,2^ This, however, does not imply that IDPs are fully random polymers. Rather, it is now acknowledged that a combination of conformational ensembles and possibly average structural properties is necessary to describe the IDP functional correlates.^3^ For instance, the ensemble averaged size of the IDPs (i.e. radius of gyration and polymer scaling exponent) was found to correlate with its tendency of liquid-liquid phase separation (LLPS) and potential for forming membraneless organelles.^4,5^ The structural property of an IDP is known to be easily affected by external conditions such as temperature,^6–9^ salt,^10–13^ pH,^14^ posttranslational modifications^15^ and the other biomolecular interaction partners (i.e. other proteins and nucleic acids). Among these, salt is one of the most frequent way to perturb IDP conformations in vitro, which has attracted both theoretical and experimental attention.^10,16–18^

Theoretically, the net charge per residue and fraction of charged amino acids of an IDP sequence can be used to understand any salt-induced conformational changes from electrostatic screening.^19^ Polyampholyte theory works well in quantitatively predicting variation in size of IDPs over different salt concentrations.^20^ A sequence whose charged amino acids are more closely balanced (close to zero net charge) would result in salt-induced expansion. However, this observation could break down for some cases when charge patterning emerges.^16,17,21^ A recent extension on the charge patterning variable, sequence charge decoration (*SCD*), at low and high salt regime (*SCD*_lowsalt_ and *SCD*_highsalt_) has been shown to provide fast, qualitative predictions of salt-induced trends.^12^ Alternatively, all-atom implicit or explicit-solvent simulations^13,22^ and coarse-grained simulations with a Debye-Hückel screening term^23,24^ can be applied to investigate the impact of electrostatic screening, albeit with more computational resources.

Beyond electrostatic screening, another important mechanism can impact the effect of salt on IDP conformations. In its simplest case, salt is known to change the solubility of amino acids, which is both amino acid and salt-type dependent.^25–27^ If the solubility of an amino acid reduces with increasing salt concentrations, this is referred to as a ‘salting-out’ effect, whereas increasing solubility with increasing salt concentrations is named ‘salting-in’. The behaviors of specific ions in salting-out/in the proteins are catalogued in the Hofmeister series.^25^ The salting-out effect is believed to affect the conformation of IDPs in the high salt regime whereas electrostatic screening dominates salt-induced conformational change in the low salt regime. The turning point between the two effects is known to be above physiological salt concentration,^13^ but may be sequence dependent and correlate with the fraction of charged amino acids. In other words, the salting-out effect could become more predominant for a sequence with limited charged amino acids. The low complexity domain of FUS protein, for instance, has only two negatively charged amino acids out of 163 and has been shown to start phase separating with increasing salt concentrations.^28^ Considering the wide variety of disordered sequences with limited charged amino acids and increasing amount of salt-dependent observations, it is necessary to introduce a computational model with both electrostatic screening and salting-out effect.

When looking at the amino acid dependent salting-out effect, all-atom explicit-solvent simulation is the most straightforward way. Due to the sampling difficulty, however, such simulations are usually unreachable for proteins with more than 100 amino acids without the use of specialized supercomputers.^29,30^ It should also be noted that it is unclear how well the current salt force field can reproduce salt-induced conformational changes of IDPs, especially at high salt concentrations.^13,31^ This is mainly due to the limited experimental data available for balancing the interactions among protein, water, and salt. The effect of the salt solution on specific amino acids can also be described by the salting-out constant,^32^ which can be considered the free energy of transferring the amino acid from water to 1M of salt solution. A salt-dependent hydropathy scale can therefore be introduced using these salting-out constants. A simple sequence descriptor which considers the hydrophobic patterning^33,34^ is the first option to implement such a scale. However, considering the challenge of balancing the contributions from charge and hydrophobic patterning,^33^ a coarse-grained model is the most cost effective way to implement the salting-out effect.

In this work, we first collect a set of 17 proteins with existing experimental observations of salt-dependent LLPS behavior and ask if electrostatic screening is sufficient to explain their behavior. We then introduce an amino-acid specific salting-out term into our coarse-grained model, optimize it with a recent Förster resonance energy transfer (FRET) data set,^21^ and verify the model with the salt-dependent LLPS data set. At last we ask how much the salting-out effect would affect the conformation of a typical IDP sequence, providing guidance for future IDP research when varying the salt conditions.

## Role of electrostatic screening

Since there are two different mechanisms by which salt could alter the conformation of an IDP (i.e. electrostatic screening and salting-out), we would like to first test how much can electrostatic screening alone explain the experimentally determined salt-dependent behavior of IDPs. Thanks to the rapidly growing field of membraneless organelles and LLPS, there have been a number of measurements of various IDPs across a range of salt concentrations. Here we start by going through all the entries in an LLPS database^35^ and selecting a total of 17 sequences with experimentally measured salt-dependent LLPS behaviors.^28,36–51^ We classify the sequences with different salt-dependent LLPS behaviors into three groups. The first group of IDP sequences do not form liquid droplets at low salt concentrations and LLPS is observed with increasing salt concentrations. This group will be referred as high-salt phase separation (HSPS). The second group of IDP sequences phase separate at low salt concentrations and the liquid droplets disappear at high salt concentrations, which we refer to as low-salt phase separation (LSPS). There could also be sequences that phase separate at medium range of salt concentrations and the droplets disappear at both low and high salt concentrations. This will be called medium-salt phase separation (MSPS). Of all the 17 sequences we have collected, 6 of them behave as HSPS, 10 as LSPS and 1 as MSPS. We organize the description of these sequences including the references and phase behaviors using different models in Table S1 and the sequences in Supporting Methods 1.1.

The next step was to ask if the existing sequence descriptor *SCD*_lowsalt_^12^ (see Supporting Methods 1.2) could capture these behaviors. *SCD*_lowsalt_ characterizes the variation of protein size at the low salt concentration limit due to charge patterning. A positive value of *SCD*_lowsalt_ suggests overall repulsive intramolecular interactions, which will be reduced with increasing salt concentrations and electrostatic screening leading to protein collapse. In order to compare directly with the experimental salt-dependent phase behaviors, we need to consider the known correlation between the protein radius of gyration (*R*_g_) and the ease of phase separation. That is, a reducing *R*_g_ (collapsing) suggests increasing amino acid interactions and consequently more prone to phase separation.^4,5^ A positive *SCD*_lowsalt_, therefore, predicts the sequence will phase separate at high salt concentrations (HSPS). Meanwhile, a negative *SCD*_lowsalt_ predicts phase separation at low salt concentrations (LSPS). As shown in Fig. 1B, 5 of the 6 proteins in the first group with experimentally observed HSPS behavior show positive *SCD*_lowsalt_. However 2 of them (i.e. FUS LC and TDP-43 CTD) have an *SCD*_lowsalt_ value close to zero (*<*0.3) due to their limited fraction of charged amino acids per residue (*f*_*q*_ *<*5%) shown in Fig. 1A. This is expected since *SCD*_lowsalt_ was designed to capture the impact of charge patterning on protein size at low salt regime and might not provide an obvious trend for sequences with low *f*_*q*_. *SCD*_lowsalt_ therefore works for correctly predicting the phase behavior of 3 of the 6 HSPS proteins. When looking at the second group of proteins with LSPS behaviors, 2 of the 10 proteins have negative *SCD*_lowsalt_ values. For the only protein in the third group, a positive *SCD*_lowsalt_ correctly captures the trend at low salt limit. For the high salt limit, however, we still see a positive *SCD*_highsalt_ value (see Supporting Methods 1.2 and Fig. S1), predicting uniform HSPS instead of MSPS observed in the experiment. Overall, the charge patterning descriptor can capture the salt-dependent phase behaviors of 5 of the 17 protein sequences.

**Figure 1:**
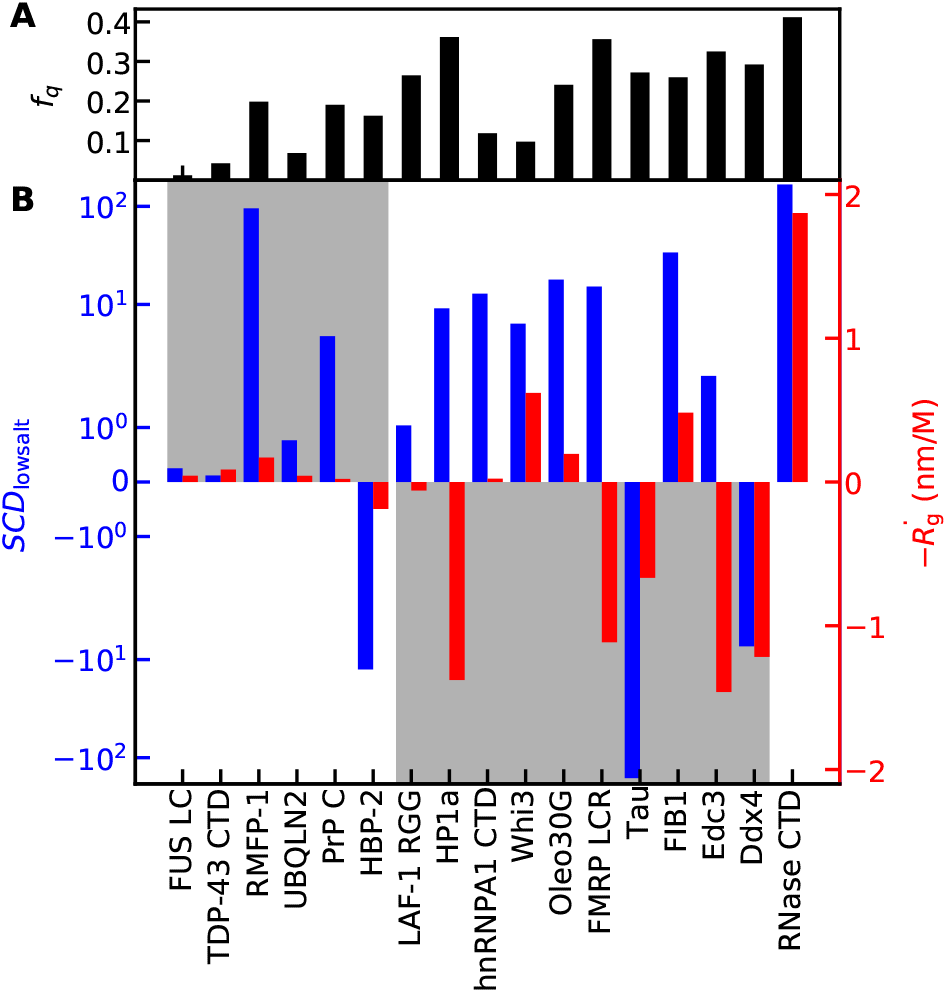
Capturing the salt-dependent LLPS behaviors with electrostatic screening. A) The fraction of charged amino acids (*f*_*q*_) for the sequences tested. B) *SCD*_lowsalt_ (blue) and the negative derivative of radius of gyration with respect to salt concentrations (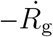, red) using the HPS model. The gray shading shows the expected behaviors on these values using experimentally observed salt-dependent phase behaviors.

To explicitly consider the impact of the electrostatic screening on IDP conformations, a coarse-grained (CG) model can be used. We simulate the *R*_g_ at a wide range of salt concentrations using the HPS model^24^ (see Supporting Methods 1.3) in which the salt concentration is captured by a Debye-Hückel screening term.^23^ In order to compare with the experimentally observed LLPS behaviors, we calculate the derivative of *R*_g_ with respect to the salt concentration 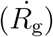 at the middle of the experimental salt range (Table S1). A negative 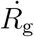 suggests a reducing *R*_g_ with increasing salt, stronger interactions, and easier phase separation at high salt concentrations (HSPS). A positive, on the other hand, 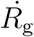 suggests an expansion of the protein and LSPS. Since a negative 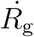 is comparable to a positive *SCD*_lowsalt_ and HSPS, we plot in Fig. 1B 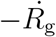 together with *SCD*_lowsalt_. We find that 4 of the 6 sequences have an almost zero 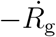 (*<*0.1 nm/M), whereas only 1 of the 6 shows a clear HSPS behavior. For the proteins with LSPS behaviors, the HPS model can capture 5 of the 10. For the only protein with the MSPS behavior, the *R*_g_ does first reduce and then increase (see RNase CTD in Fig. S2). The CG model is then consistent with the experimental observation of appearing and disappearing liquid droplets of RNase CTD.^51^ Altogether, our CG model captures the salt-dependent phase behaviors of 6 of the 17 protein sequences, with almost negligible improvements in comparing to *SCD*, suggesting a mechanism in addition to electrostatic screening should be taken into account.

## HPS model with the salting-out effect

The effectiveness of salt on amino acids can be quantified by the Setschenow equation^52^

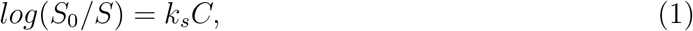

in which *S*_0_ and *S* are the solubility of amino acids at water and salt conditions, respectively, and *k*_*s*_ is the salting-out constant. A positive *k*_*s*_ value suggests that the amino acid is less soluble with increasing salt concentrations. This is commonly referred to as ‘salting-out’. Meanwhile, a negative *k*_*s*_ value suggests the amino acid solubility increases with increasing salt concentrations, known as ‘salting-in’. For different type of salts, the salting-out effect on the same amino acid could vary, which is described by the Hofmeister series.^25^ Here we would like to focus on providing a model for NaCl which is commonly used in the experiment to perturb IDP conformations. The salting-out constants for half of the twenty amino acids can be found in old literature,^26,32,53,54^ whereas the rest can be obtained by using the empirical correlation between the salting-out constants and the amino acid hydropathy scales^55^ as shown previously^24^ and in Fig. 2A.

**Figure 2:**
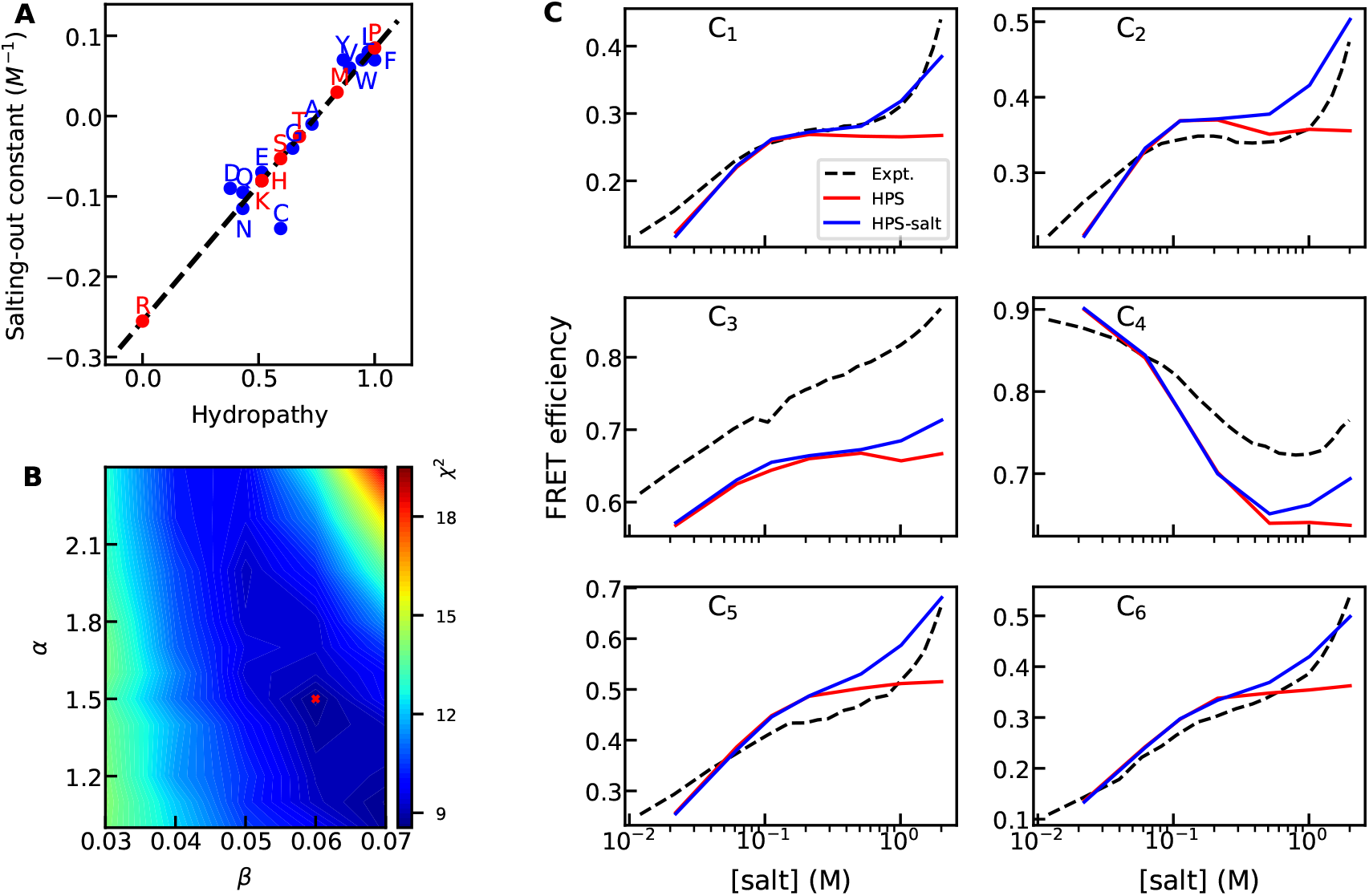
A) The correlation between salting-out constants and hydropathy scales. Black line shows the linear fitting curve between these two parameters. Blue dots show the data from literature^26,32,53,54^ and red dots show the estimate from linear interpolation or extrapolation. The numerical values of these salting-out coefficients are shown in Table S2. B) The deviation *χ*^2^ from the experimental data for using different sets of free parameters in the salting-out term. The red cross shows the combination of the two parameters with the smallest *χ*^2^. C) FRET efficiencies of six constructs (labeling positions) of E-cad from the experiment, HPS model and HPS-salt model using the best combination of free parameters shown in B.

We can therefore introduce a linear adjustment to the amino acid hydropathy scales based on the salting-out constants *k*_*s*_. The new salt-dependent hydropathy in the HPS model can be written as

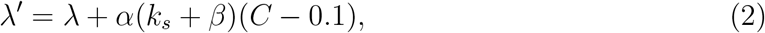

in which *C* is the salt concentration and *α* and *β* are the two free parameters. *α* controls how large a concentration-dependent correction we need for the amino acid hydropathy whereas *β* tunes the behavior of amino acids with close to zero salting-out constants. In order to optimize these two parameters, experimental measurements of IDP size at different salt concentrations, for example from FRET or small-angle X-ray scattering (SAXS), should be the most straightforward target data. However since electrostatic screening usually dominates the size variation at low salt concentrations and salting-out effect would only become more prominent beyond 0.5M ionic strength,^13^ very limited experimental data is available.

A most recent set of FRET measurements of the disordered tail of the cell-adhesion protein E-cadherin^56,57^ (hereafter, E-cad) provides a unique data set for us to optimize the two free parameters for this salting-out term.^21^ The experimental data set contains six different pair labeling positions along E-cad spanning a wide range of salt concentrations. As shown in black dashed lines of Fig. 2C, the salt dependent conformation of E-cad differs for the six constructs: five of the six constructs always collapse with increasing salt concentrations whereas one of them (C_4_) first expands and then starts to collapse at about 0.5M salt concentration. In the same work, we have tried to use the HPS model to capture the salt-dependent conformational change of E-cad for these six constructs at low salt concentrations and were able to reproduce the different behavior of C_4_ in contrast to the other five constructs.^21^ The reason for the unique salt-dependent trend of C_4_ from the other constructs is due to the balanced positively and negatively charged amino acids in C_4_. This fragment behaves more like a polyampholyte,^20^ whereas the other constructs with unbalanced charges are closer to a polyelectrolyte and collapse with increasing electrostatic screening. What we could not capture in that work was the salt dependent behaviors above 0.2M. In addition to C_4_, all the other constructs also show a different tendency of collapsing above 0.2M, which cannot be captured by the HPS model. We believe this is due to the absence of salting-out effect in the HPS model.

We therefore scan a range of the two free parameters and calculate the deviations of the simulated FRET efficiencies from the experimental measurements. As shown in Fig. 2B, we show the *χ*^2^ defined as 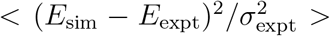 from all the six constructs, in which *σ*_expt_ (=0.03) is the experimental error of FRET efficiencies. We find the best combination of parameters to be *α*=1.5 and *β*=0.06. We note a nonzero *β* value suggests a necessary shift for the current set of salting-out constants so that some of the amino acids with values close to zero might change their behavior from salting-in to salting-out. This could result from two sources of errors. First, salting-out constants of amino acids were usually estimated by summing up constants from small functional groups with the assumption that the effect is additive.^32^ Second, a different hydropathy scale can be used for extrapolating a different version of salting-out constants. There have been multiple recent works on optimizing the hydropathy scale for the HPS model based on reproducing the experimental radii of gyration and/or the LLPS behaviors of IDPs at physiological salt concentrations.^58–60^ However, the limited salt-dependent experimental data set available does not allow us to address further these two sources of errors. With the best set of *α* and *β*, we show in Fig. 2C that the different trends seen above 0.2M can now be captured. We refer to the new model as the HPS-salt model.

## Salt-dependent liquid-liquid phase separation

To verify the HPS-salt model, we apply it to the 17 sequences (see Table S1) with experimentally accessible salt-dependent data and compare the results with the HPS model (see Supporting Methods 1.3). We use the interaction strength *ϵ* (0.2 kcal/mol) of the HPS model, but also provide the results of the 17 sequences using a different *ϵ* (0.16 kcal/mol) in Fig. S3. We find that all the salt-dependent results discussed below are not affected by small variations of *ϵ* since it changes the *R*_g_ obtained from the model at all salt concentrations similarly.

We first look at the sequences with HSPS behaviors shown in Fig. 3A. We find that the *R*_g_ for all these sequences decrease with rising salt concentration, suggesting increasing interactions between amino acids. Based on the correlation between single-chain properties like *R*_g_ and LLPS behaviors,^4,5^ these sequences are predicted to phase separate more readily at high salt concentration, consistent with the experimental observations. On the other hand, the original HPS model without the salting-out term can only capture the HSPS behaviors of 1 of the 6 sequences. To understand this improvement, we calculate the sequence hydropathy decoration (*SHD*), which has recently been shown to quantitatively describe the hydropathy patterning of the sequence.^33^ *SHD* takes the hydropathy scale of the amino acids along the sequence together with their sequence separation to estimate the size of the IDP, similar to what *SCD* does for considering the contribution of charged amino acids.^17^ A salt-dependent variant of *SHD* (*SHD*_salt_, see Supporting Methods 1.2) can be defined in which *λ* is replaced with salt-dependent *λ*′ using the same optimized parameters in the previous section (Eq. 2). We plot *SHD*_salt_ as a function of the salt concentrations for these sequences in Fig. 3D. It is clear that all these sequences have an increasing *SHD*_salt_ with increasing salt concentrations, suggesting stronger interactions and therefore ease of LLPS. It is worth noting that of the 5 sequences with improving predictions, only 2 of them feature a low fraction of charged amino acids (*<* 5%, see Fig. 1A). This suggests that the salting out effect, which was considered less important than electrostatic screening at low salt concentrations, can also play a role for sequences with decent fraction of charged amino acids.

**Figure 3:**
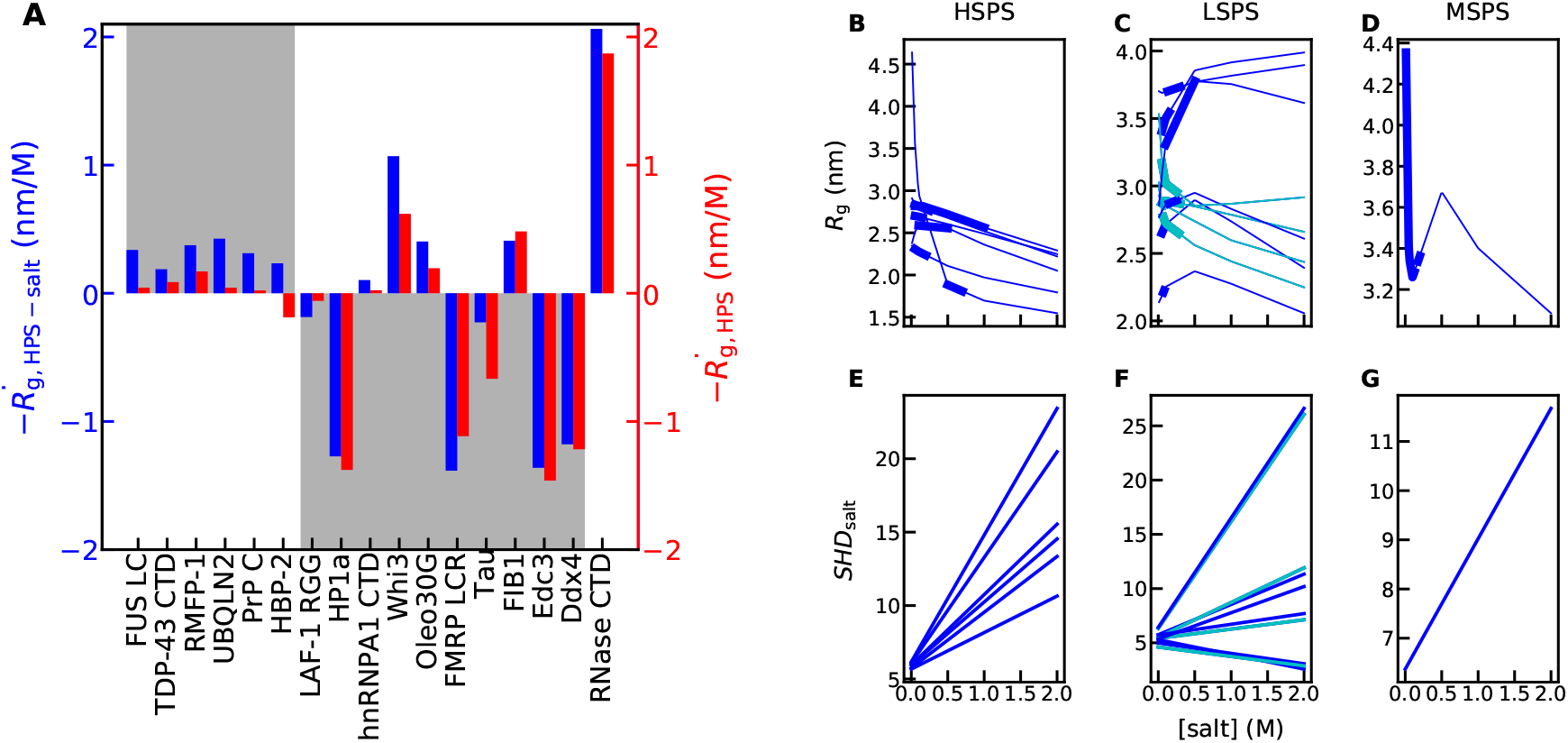
Salt-dependent LLPS using the HPS-salt model. A) Salt-dependent trend of *R*_g_ (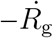 obtained using the HPS-salt and HPS models. B, C and D) Salt-dependent *R*_*g*_ of the 17 sequences using the HPS-salt model grouped into three clusters with experimentally-observed HSPS, LSPS and MSPS behaviors, respectively. Thick lines show the range of experimental salt concentrations. Blue lines show the sequences the salt-dependent LLPS behaviors of which can be captured by the model whereas the cyan lines show the sequences with behaviors that the HPS-salt model can not capture. E, F and G) The corresponding *SHD*_salt_ as a function of salt concentrations.

For the sequences with experimentally observed LSPS behaviors, the original HPS model can capture 5 of the 10. The HPS-salt model is able to capture one additional sequence, LAF-1 RGG (Fig. 3B), which is due to a reduced *SHD*_salt_ at higher salt concentrations. However, there are still four sequences (cyan in Fig. 3B and E) for which the HPS-salt model still predicts a decreasing *R*_g_ with rising salt concentrations: hnRNPA1 CTD, Whi3, Oleo30G, and FIB1. Only 1 out of these 4 sequences has a decreasing *SHD*_salt_ when increasing salinity which might help with the correction. There could be a few causes for such deviations. First, the current salting-out term is parameterized using only the FRET data of E-cad and might not be applicable to these four sequences. Second, the hydropathy used in the current HPS model might require further optimization as suggested in recent publications.^58–60^ Third, there may be local or long-range interactions that our simple CG model cannot capture. For instance, Oleo30G has a short fragment with high helical propensity in the middle, which is not considered in the current model.^45^

For the only sequence with experimentally observed MSPS behavior, both the HPS-salt and HPS models can capture that. These suggest that for sequences with LSPS and MSPS behaviors, the additional salting-out term only provides slight improvement against the HPS model. Comparing with the sequence descriptor *SCD*_lowsalt_, most of the improvements for LSPS sequences come from CG simulations instead of the salting-out term.

Finally, we would like to investigate how the salting out effect applies to a wide variety of disordered proteins. We employ sequences from the Disprot database.^61,62^ For each sequence, only the longest disordered region is selected. We exclude sequences with a disordered region shorter than 30 residues, as the polymer theory may not work well for shorter chain lengths, and longer than 400 residues as sampling becomes more difficult for longer sequences using CG simulations. We simulate the *R*_g_ for a total of 530 sequences at different ionic strengths and compare both *R*_g_ and 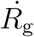 (i.e. the derivative of *R*_g_ along the salt concentration) obtained using the HPS and HPS-salt models. As shown in Fig. 4A, the differences of *R*_g_ between the two models (Δ*R*_g_) are exactly the same for all the sequences at 0.1M because the additive salting-out term is designed to be zero at that condition. As expected when increasing the ionic strength, more deviations between the two models are observed since the salting-out term is linearly dependent on the salt concentration (Fig. 4B and C).

**Figure 4:**
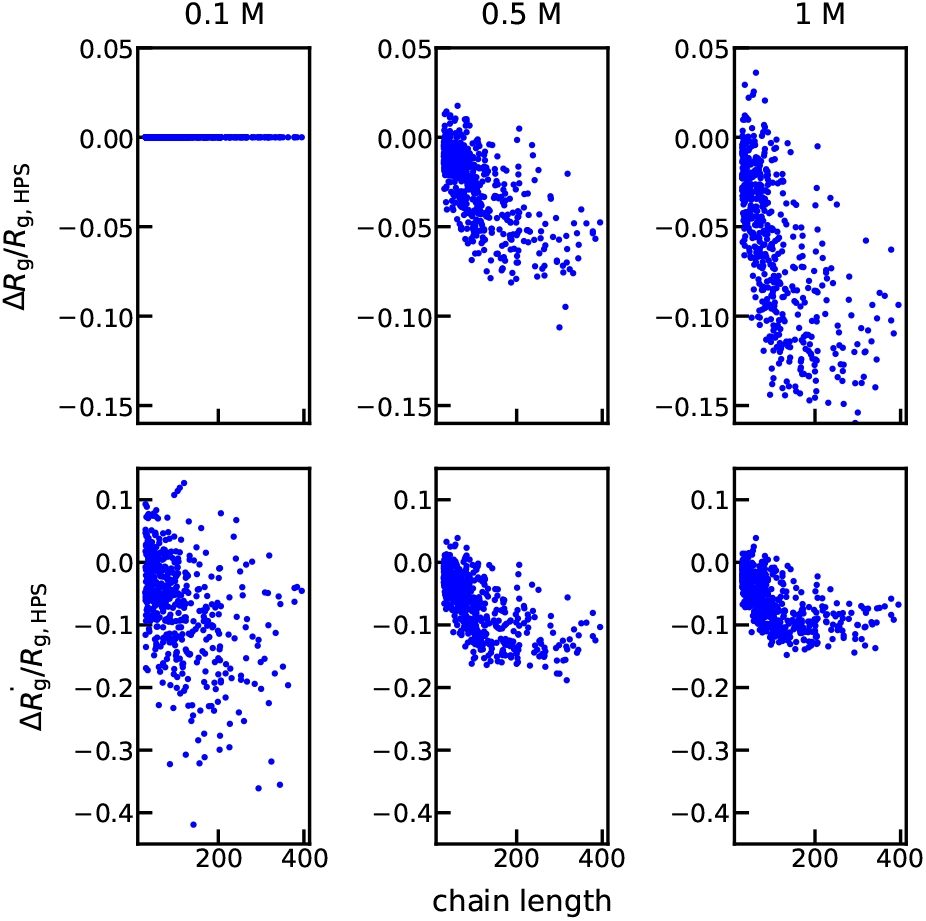
Top: Difference between *R*_g_ from HPS-salt and HPS models at different salt concentrations for the IDP sequences from the Disprot database.^61,62^ Bottom: Difference between slopes of *R*_g_ from the HPS-salt and HPS models. All values are normalized by the *R*_g_ from the HPS model at the corresponding salt concentration.

Interestingly, the difference between 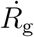 of the two models 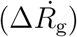 (Fig. 4D, E and F) does not behave the same as Δ*R*_g_. If we only look at the consistency between the signs of 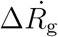 from the two models, about 14%, 36% and 50% of the sequences show different signs at 0.1M, 0.5M and 1.0M, respectively (see Fig. S4). This is consistent with the observation of Δ*R*_g_, in which the salting-out effect is more significant in correcting *R*_g_ predictions at high salt concentrations. However for individual sequences, large 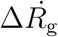 appear more frequently at low salt concentrations (bottom panels of Fig. 4). This is due to the fact that electrostatic screening only introduces size variation at low salt concentrations, and introducing the salting-out effect is more likely to create variations in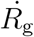 as long as the effect of electrostatic screening and salting-out have opposite signs. Such a counter-intuitive observation suggests that even though the salting-out effect might be secondary to electrostatic screening when considering general IDP sequences at physiological salt concentrations, for some IDP sequences an incorrect prediction of salt dependence can be easily introduced if the salting-out effect is not considered. We have also checked if 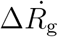 correlates with fraction of charged amino acids (*f*_c_) and find no direct evidence (see Fig. S5). This is because the role of electrostatic screening becomes important only when the sequence contains reasonable amount of charged amino acids and the salting-out effect can then introduce large salt-dependent deviations from electrostatic screening.

## Conclusion

In this work, we introduced a method to incorporate the salting-out effect into the coarse-grained HPS model. The new HPS-salt model used the experimentally measured salting-out constants together with the FRET measurement of an IDP to parameterize a salting-out term in the HPS model. We further verified the model by collecting 17 disordered sequences with experimentally observed salt-dependent LLPS behaviors. This entire scheme of introducing a salt-dependent term into the hydropathy used in the HPS model can also benefit from the recent efforts on improving the hydropathy scale^58–60^ and can be easily extended to other simulation models.

We find that the new HPS-salt model improved the predictions by capturing the salt-dependent LLPS behaviors of 13 of the 17 sequences rather than 6 of the 17 sequences using the HPS model. This suggests electrostatic screening alone is not sufficient for some IDP sequences, not just limited to the ones with few charged amino acids. We also assessed in general the roles of the salting-out term and electrostatic screening in varying salt-dependent conformations of IDPs. Interestingly, even though the salting-out effect mostly affects the salt-dependent trend of radius of gyration at high salt concentrations, its impact to radius of gyration can be completely opposite to electrostatic screening. This results in larger deviations between the two mechanisms at low salt concentrations, where electrostatic screening was thought to dominate, than those at high salt concentration. We therefore conclude the salting-out effect, which was usually considered to be a secondary effect to the electrostatic screening, is not limited to IDP sequences with low fraction of charged amino acids and can also be important in predicting the salt-dependent IDP conformations at low salt concentrations.

## Acknowledgement

This work was supported by the National Science Foundation grant MCB-2015030 and the National Institutes of Health grant R01GM120537. The authors acknowledge Research Computing at Arizona State University for providing HPC and storage resources.

## References

(1) Uversky, V. N.; Gillespie, J. R.; Fink, A. L. Why are “natively unfolded” proteins unstructured under physiologic conditions? Proteins 2000, 41, 415–427.

(2) Wright, P. E.; Dyson, H. J. Intrinsically disordered proteins in cellular signalling and regulation. Nat. Rev. Mol. Cell Biol. 2015, 16, 18–29.

(3) Fisher, C. K.; Stultz, C. M. Constructing ensembles for intrinsically disordered proteins. Curr. Opin. Struct. Biol. 2011, 21, 426–431.

(4) Lin, Y.-H.; Chan, H. S. Phase Separation and Single-Chain Compactness of Charged Disordered Proteins Are Strongly Correlated. Biophys. J. 2017, 112, 2043–2046.

(5) Dignon, G. L.; Zheng, W.; Best, R. B.; Kim, Y. C.; Mittal, J. Relation between single-molecule properties and phase behavior of intrinsically disordered proteins. Proc. Natl. Acad. Sci. U.S.A. 2018, 115, 9929–9934.

(6) Wuttke, R.; Hofmann, H.; Nettels, D.; Borgia, M. B.; Mittal, J.; Best, R. B.; Schuler, B. Temperature-dependent solvation modulates the dimensions of disordered proteins. Proc. Natl. Acad. Sci. U.S.A. 2014, 111, 5213–5218.

(7) Zerze, G. H.; Best, R. B.; Mittal, J. Sequence-and temperature-dependent properties of unfolded and disordered proteins from atomistic simulations. J. Phys. Chem. B 2015, 119, 14622–14630.

(8) Dignon, G. L.; Zheng, W.; Kim, Y. C.; Mittal, J. Temperature-Controlled Liquid– Liquid Phase Separation of Disordered Proteins. ACS Cent. Sci. 2019, 5, 821.

(9) Dong, X.; Bera, S.; Qiao, Q.; Tang, Y.; Lao, Z.; Luo, Y.; Gazit, E.; Wei, G. Liquid– Liquid Phase Separation of Tau Protein Is Encoded at the Monomeric Level. J. Phys. Chem. Lett. 2021, 12, 2576–2586.

(10) Muller-Spath, S.; Sorrano, A.; Hirschfeld, V.; Hofmann, H.; Ruegger, S.; Reymond, L.; Nettels, D.; Schuler, B. Charge interactions can dominate the dimensions of intrinsically disordered proteins. Proc. Natl. Acad. Sci. U.S.A. 2010, 107, 14609–14614.

(11) Borgia, A.; Borgia, M. B.; Bugge, K.; Kissling, V. M.; Heidarsson, P. O.; Fernandes, C. B.; Sottini, A.; Buholzer, K. J.; Nettels, D.; Kragelund, B. B. et al. Extreme disorder in an ultra-high-affinity protein complex. Nature 2018, 555, 61–66.

(12) Huihui, J.; Firman, T.; Ghosh, K. Modulating charge patterning and ionic strength as a strategy to induce conformational changes in intrinsically disordered proteins. J. Chem. Phys. 2018, 149, 085101.

(13) Vancraenenbroeck, R.; Harel, Y. S.; Zheng, W.; Hofmann, H. Polymer effects modulate binding affinities in disordered proteins. Proc. Natl. Acad. Sci. U.S.A. 2019, 116, 19506–19512.

(14) Kjaergaard, M.; Brander, S.; Poulsen, F. M. Random coil chemical shift for intrinsically disordered proteins: effects of temperature and pH. J. Biomol. NMR 2011, 49, 139–149.

(15) Martin, E. W.; Holehouse, A. S.; Grace, C. R.; Hughes, A. J.; Pappu, R. V. Sequence determinants of the conformational properties of an intrinsically disordered protein prior to and upon multisite phosphorylation. J. Am. Chem. Soc. 2016, 138, 15323– 15335.

(16) Das, R. K.; Pappu, R. V. Conformations of intrinsically disordered proteins are influenced by linear sequence distributions of oppositely charged residues. Proc. Natl. Acad. Sci. U.S.A. 2013, 110, 13392–13397.

(17) Sawle, L.; Ghosh, K. A theoretical method to compute sequence dependent configurational properties in charged polymers and proteins. J. Chem. Phys. 2015, 143, 085101.

(18) Samanta, H. S.; Chakraborty, D.; Thirumalai, D. Charge fluctuation effects on the shape of flexible polyampholytes with applications to intrinsically disordered proteins. J. Chem. Phys. 2018, 149, 163323.

(19) Mao, A. H.; Crick, S. L.; Vitalis, A.; Chicoine, C.; Pappu, R. V. Net charge per residue modulates conformational ensembles of intriniscally disordered proteins. Proc. Natl. Acad. Sci. U.S.A. 2010, 107, 8183–8188.

(20) Higgs, P. G.; Joanny, J.-F. Theory of polyampholyte solutions. J. Chem. Phys. 1991, 94, 1543–1554.

(21) Wiggers, F.; Wohl, S.; Dubovetskyi, A.; Rosenblum, G.; Zheng, W.; Hofmann, H. Diffusion of the disordered E-cadherin tail on β-catenin. bioRxiv 2021, https://www.biorxiv.org/content/10.1101/2021.02.03.429507v1.

(22) Vitalis, A.; Pappu, R. V. ABSINTH: A new continuum solvent model for simulations of polypeptides in aqueous solutions. J. Comput. Chem. 2008, 30, 673–699.

(23) Debye, P.; Huckel, E. De la theorie des electrolytes. I. abaissement du point de congelation et phenomenes associes. Physikalische Zeitschrift 1923, 24, 185–206.

(24) Dignon, G. L.; Zheng, W.; Kim, Y. C.; Best, R. B.; Mittal, J. Sequence determinants of protein phase behavior from a coarse-grained model. PLoS Comput. Biol. 2018, 14, e1005941.

(25) Hofmeister, F. Zur lehre von der wirkung der salze. Archiv fur experimentelle Pathologie und Pharmakologie 1888, 24, 247–260.

(26) Baldwin, R. L. How Hofmeister ion interactions affect protein stability. Biophys. J. 1996, 71, 2056–2063.

(27) Zhou, H.-X. Interactions of macromolecules with salt ions: an electrostatic theory for the Hofmeister effect. Proteins 2005, 61, 69–78.

(28) Burke, K. A.; Janke, A. M.; Rhine, C. L.; Fawzi, N. L. Residue-by-residue view of in vitro FUS granules that bind the C-terminal domain of RNA polymerase II. Mol. Cell 2015, 60, 231–241.

(29) Shaw, D. E.; Grossman, J.; Bank, J. A.; Batson, B.; Butts, J. A.; Chao, J. C.; Deneroff, M. M.; Dror, R. O.; Even, A.; Fenton, C. H. et al. Anton 2: raising the bar for performance and programmability in a special-purpose molecular dynamics super-computer. Proceedings of the international conference for high performance computing, networking, storage and analysis. 2014; pp 41–53.

(30) Zheng, W.; Dignon, G. L.; Jovic, N.; Xu, X.; Regy, R. M.; Fawzi, N. L.; Kim, Y. C.; Best, R. B.; Mittal, J. Molecular Details of Protein Condensates Probed by Microsecond Long Atomistic Simulations. J. Phys. Chem. B 2020, 124, 11671—-11679.

(31) Luo, Y.; Roux, B. Simulation of osmotic pressure in concentrated aqueous salt solutions. J. Phys. Chem. Lett. 2010, 1, 183–189.

(32) Nandi, P. K.; Robinson, D. R. Effects of salts on the free energy of the peptide group. J. Am. Chem. Soc. 1972, 94, 1299–1308.

(33) Zheng, W.; Dignon, G.; Brown, M.; Kim, Y. C.; Mittal, J. Hydropathy patterning complements charge patterning to describe conformational preferences of disordered proteins. J. Phys. Chem. Lett. 2020, 11, 3408–3415.

(34) Bowman, M. A.; Riback, J. A.; Rodriguez, A.; Guo, H.; Li, J.; Sosnick, T. R.; Clark, P. L. Properties of protein unfolded states suggest broad selection for expanded conformational ensembles. Proc. Natl. Acad. Sci. U.S.A. 2020, 117, 23356–23364.

(35) Li, Q.; Peng, X.; Li, Y.; Tang, W.; Zhu, J.; Huang, J.; Qi, Y.; Zhang, Z. LLPSDB: a database of proteins undergoing liquid–liquid phase separation in vitro. Nucleic Acids Res. 2020, 48, D320–D327.

(36) Conicella, A. E.; Zerze, G. H.; Mittal, J.; Fawzi, N. L. ALS mutations disrupt phase separation mediated by α-helical structure in the TDP-43 low-complexity C-terminal domain. Structure 2016, 24, 1537–1549.

(37) Kim, S.; Yoo, H. Y.; Huang, J.; Lee, Y.; Park, S.; Park, Y.; Jin, S.; Jung, Y. M.; Zeng, H.; Hwang, D. S. et al. Salt triggers the simple coacervation of an underwater adhesive when cations meet aromatic π electrons in seawater. ACS Nano 2017, 11, 6764–6772.

(38) Yang, Y.; Jones, H. B.; Dao, T. P.; Castañeda, C. A. Single amino acid substitutions in stickers, but not spacers, substantially alter UBQLN2 phase transitions and dense phase material properties. J. Phys. Chem. B 2019, 123, 3618–3629.

(39) Kostylev, M. A.; Tuttle, M. D.; Lee, S.; Klein, L. E.; Takahashi, H.; Cox, T. O.; Gunther, E. C.; Zilm, K. W.; Strittmatter, S. M. Liquid and hydrogel phases of PrPC linked to conformation shifts and triggered by alzheimer’s amyloid-β oligomers. Mol. Cell 2018, 72, 426–443.

(40) Le Ferrand, H.; Duchamp, M.; Gabryelczyk, B.; Cai, H.; Miserez, A. Time-resolved observations of liquid–liquid phase separation at the nanoscale using in situ liquid transmission electron microscopy. J. Am. Chem. Soc. 2019, 141, 7202–7210.

(41) Elbaum-Garfinkle, S.; Kim, Y.; Szczepaniak, K.; Chen, C. C.-H.; Eckmann, C. R.; Myong, S.; Brangwynne, C. P. The disordered P granule protein LAF-1 drives phase separation into droplets with tunable viscosity and dynamics. Proc. Natl. Acad. Sci. U.S.A. 2015, 112, 7189–7194.

(42) Strom, A. R.; Emelyanov, A. V.; Mir, M.; Fyodorov, D. V.; Darzacq, X.; Karpen, G. H. Phase separation drives heterochromatin domain formation. Nature 2017, 547, 241– 245.

(43) Martin, E. W.; Thomasen, F. E.; Milkovic, N. M.; Cuneo, M. J.; Grace, C. R.; Nourse, A.; Lindorff-Larsen, K.; Mittag, T. Interplay of folded domains and the disordered low-complexity domain in mediating hnRNPA1 phase separation. Nucleic Acids Res. 2021, 49, 2931–2945.

(44) Zhang, H.; Elbaum-Garfinkle, S.; Langdon, E. M.; Taylor, N.; Occhipinti, P.; Bridges, A. A.; Brangwynne, C. P.; Gladfelter, A. S. RNA controls PolyQ protein phase transitions. Mol. Cell 2015, 60, 220–230.

(45) Reed, E. H.; Hammer, D. A. Redox sensitive protein droplets from recombinant oleosin. Soft Matter 2018, 14, 6506–6513.

(46) Tsang, B.; Arsenault, J.; Vernon, R. M.; Lin, H.; Sonenberg, N.; Wang, L.-Y.; Bah, A.; Forman-Kay, J. D. Phosphoregulated FMRP phase separation models activity-dependent translation through bidirectional control of mRNA granule formation. Proc. Natl. Acad. Sci. U.S.A. 2019, 116, 4218–4227.

(47) Hernández-Vega, A.; Braun, M.; Scharrel, L.; Jahnel, M.; Wegmann, S.; Hyman, B. T.; Alberti, S.; Diez, S.; Hyman, A. A. Local nucleation of microtubule bundles through tubulin concentration into a condensed tau phase. Cell Rep. 2017, 20, 2304–2312.

(48) Feric, M.; Vaidya, N.; Harmon, T. S.; Mitrea, D. M.; Zhu, L.; Richardson, T. M.; Kriwacki, R. W.; Pappu, R. V.; Brangwynne, C. P. Coexisting liquid phases underlie nucleolar subcompartments. Cell 2016, 165, 1686–1697.

(49) Schutz, S.; Noldeke, E. R.; Sprangers, R. A synergistic network of interactions promotes the formation of in vitro processing bodies and protects mRNA against decapping. Nucleic Acids Res. 2017, 45, 6911–6922.

(50) Brady, J. P.; Farber, P. J.; Sekhar, A.; Lin, Y.-H.; Huang, R.; Bah, A.; Nott, T. J.; Chan, H. S.; Baldwin, A. J.; Forman-Kay, J. D. et al. Structural and hydrodynamic properties of an intrinsically disordered region of a germ cell-specific protein on phase separation. Proc. Natl. Acad. Sci. U.S.A. 2017, 114, E8194–E8203.

(51) Al-Husini, N.; Tomares, D. T.; Bitar, O.; Childers, W. S.; Schrader, J. M. α-Proteobacterial RNA degradosomes assemble liquid-liquid phase-separated RNP bodies. Mol. Cell 2018, 71, 1027–1039.

(52) Setschenow, J. U ber die konstitution der salzlosungen auf grund ihres verhaltens zu kohlensaure. Zeitschrift fur Physikalische Chemie 1889, 4, 117–125.

(53) Schrier, E. E.; Schrier, E. B. The salting-out behavior of amides and its relation to the denaturation of proteins by salts. J. Phys. Chem. 1967, 71, 1851–1860.

(54) Nandi, P. K.; Robinson, D. R. Effects of salts on the free energies of nonpolar groups in model peptides. J. Am. Chem. Soc. 1972, 94, 1308–1315.

(55) Kapcha, L. H.; Rossky, P. J. A simple atomic-level hydrophobicity scale reveals protein interfacial structure. J. Mol. Biol. 2014, 426, 484–498.

(56) Aberle, H.; Butz, S.; Stappert, J.; Weissig, H.; Kemler, R.; Hoschuetzky, H. Assembly of the cadherin-catenin complex in vitro with recombinant proteins. J. Cell Sci. 1994, 107, 3655–3663.

(57) Huber, A. H.; Weis, W. I. The structure of the β-catenin/E-cadherin complex and the molecular basis of diverse ligand recognition by β-catenin. Cell 2001, 105, 391–402.

(58) Latham, A. P.; Zhang, B. Maximum entropy optimized force field for intrinsically disordered proteins. J. Chem. Theory Comput. 2019, 16, 773–781.

(59) Dannenhoffer-Lafage, T.; Best, R. B. A Data-Driven Hydrophobicity Scale for Predicting Liquid–Liquid Phase Separation of Proteins. J. Phys. Chem. B 2021, 125, 4046–4056.

(60) Regy, R. M.; Thompson, J.; Kim, Y. C.; Mittal, J. Improved coarse-grained model for studying sequence dependent phase separation of disordered proteins. Protein Sci. 2021, https://doi.org/10.1002/pro.4094.

(61) Sickmeier, M.; Hamilton, J. A.; LeGall, T.; Vacic, V.; Cortese, M. S.; Tantos, A.; Szabo, B.; Tompa, P.; Chen, J.; Uversky, V. N. et al. DisProt: the database of disordered proteins. Nucleic Acids Res. 2006, 35, D786–D793.

(62) Piovesan, D.; Tabaro, F.; Mičetić, I.; Necci, M.; Quaglia, F.; Oldfield, C. J.; Aspromonte, M. C.; Davey, N. E.; Davidović, R.; Dosztányi, Z. et al. DisProt 7.0: a major update of the database of disordered proteins. Nucleic Acids Res. 2017, 45, D219–D227.

