## Supplementary methods, tables and figures. for "Salt-dependent conformational changes of intrinsically disordered proteins"

### Supporting Information for: "Salt-dependent conformational changes of intrinsically disordered proteins"

### 1. Supplementary Methods

1.1. **List of protein sequences used.** Please see Table S1 for protein descriptions and references.

FUS LC
MASNDYTQQA TQSYGAYPTQ PGQGYSQQSS QPYGQQSYSG YSQSTDTSGY GQSSYSSYGQ SQNTGYGTQS TPQGYGSTGG YGSSQSSQSS YGQQSSYPGY GQQPAPSSTS GSYGSSSQSS SYGQPQSGSY SQQPSYGGQQ QSYGQQQSYN PPQGYGQQNQ YNS

TDP–43 CTD
GRFGGNPGGF GNQGGFGNSR GGGAGLGNNQ GSNMGGGMNF GAFSINPAMM AAAQAALQSS WGMMGMLASQ QNQSGPSGNN QNQGNMQREP NQAFGSGNNS YSGSNSGAAI GWGSASNAGS GSGFNGGFGS SMDSKSSGWG M

RMFP–1
M AKPSYPPTYK AKPSYPPTYK AKPSYPPTYK AKPSYPPTYK AKPSYPPTYK AKPSYPPTYK AKPSYPPTYK AKPSYPPTYK AKPSYPPTYK AKPSYPPTYK AKPSYPPTYK AKPSYPPTYK

UBQLN2
RAMQALMQIQ QGLQTLATEA PGLIPSFTPG VGVGVLGTAI GPVGPVTPIG PIGPIVPFTP IGPIGPIGPT GPAAPPGSTG SGGPTGPTVS SAAPSETTSP TSESGPNQQF IQQMVQALAG ANAPQLPNPE VRFQQQLEQL NAMGFLNREA NLQALIATGG DINAAIERLL GSQPS

PrP C
GSKKRPKPGG WNTGGSRYPG QGSPGGNRYP PQGGGGWGQP HGGGWGQPHG GGWGQPHGGG WGQPHGGGWG QGGGTHSQWN KPSKPKTNMK HMAGAAAAGA VVGGLGGYML GSAMSRPIIH FGSDYEDRYY RENMHRYPNQ VYYRPMDEYS NQNNFVHDCV NITIKQHTVT TTTKGENFTE TDVKMMERVV EQMCITQYER ESQAYYQRGS

HBP–2
MQFFGAGPFN TAHHSAVSDA AAAHHDAAGE YAQNAATGLL DTHHNENHDM THDLANGYGL HEHDEQHHGL ADGLHQEYAA RAAQGANAVH NDAAQSHSAL AAANTFGHGH APYAAYGHGV YGHGPYGHGP YGHGLYGHGL YGHGPYGHGL YGHGAFGHGL NAYAPLVGHG LRGYL

LAF–1 RGG
MESNQSNNGG SGNAALNRGG RYVPPHLRGG DGGAAAAASA GGDDRRGGAG GGGYRRGGGN SGGGGGGGYD RGYNDNRDDR DNRGGSGGYG RDRNYEDRGY NGGGGGGGNR GYNNNRGGGG GGYNRQDRGD GGSSNFSRGG YNNRDEGSDN RGSGRSYNND RRDNGGDG

HP1a

MGKKIDNPES SAKVSDAEEE EEEYAVEKII DRRVRKGKVE YYLKWKGYPE TENTWEPENN
LDCQDLIQQY EASRKDEEKS AASKKDRPSS SAKAKETQGR ASSSTSTASK RKSEEPTAPS
GNKSKRTTDA EQDTIPVSGS TGFDRGLEAE KILGASDNNG RLTFLIQFKG VDQAEMVPSS
VANEKIPRMV IHFYEERLSW YSDNED

hnRNPA1 CTD

MASASSSQRG RSGSGNFGGG RGGGFGGNDN FGRGGNFSGR GGFGGSRGGG GYGGSGDGYN
GFGNDGSNFG GGGSYNDFGN YNNQSSNFGP MKGGNFGGRS SGPYGGGGQY FAKPRNQGGY
GGSSSSSSYG SGRRF

Whi3

ELTQQQIPFQ TQMQMHQGSP PAPTHVTAYQ QPLLSASGVV SPPQSASSVK RPSLLVQRSR
FSFTDPFSSE QTNMGSQQPD LITTPLKGHQ DTGKSFLLME SDEINDSIWG NGTGIPSSIS
GLTTSQPPTP HLEWGTTGRR QSSTFYPSQS NTEIPPMHLT GQVQSSQLAT GLQQPLPQPQ
RQSLSYNLVT PLSSDMNLPP QSSQGGILPH QAPAQTQPQS QALQHHQHLH HQQQQLQQQQ
HHLQQQQHQQ QQQSLSQQPQ QQQSQQSQAH SQQHQQQHQQ QQQQQQPQQQ QPQQHPPQQP
QQQNSQQAIV GQSQQQVTSG QQKGSSRNSI SKTLQVNGPK NAAAALQNTN GISQVDLSLL
AKVPPPANPA

Oleo30G

GSTTTYDRHH VTTTQPQYRH DQHTGDRLTH PQRQQQGPST GKLALGATPL FGVIGFSPVI
VPAMGIAIGL AGVTGFQRDY VKGKLQDVGE YTGQKTKDLG QKIQHTAHEM GDQGQGQGQG
GGKEGRKEGG KLEHHHHHH

FMRP LCR

GASSRPPPNR TDKEKSYVTD DGQGMGRGSR PYRNRGHGRR GPGYTSGTNS EASNASETES
DHRDELSDWS LAPTEEERES FLRRGDGRRR GGGGRGQGGR GRGGGFKGND DHSRTDNRPR
NPREAKGRTT DGSLQIRVDC NNERSVHTKT LQNTSSEGSR LRTGKDRNQK KEKPDSVDGQ
QPLVNGVP

Tau

MAEPRQEFEV MEDHAGTYGL GDRKDQGGYT MHQDQEGDTD AGLKESPLQT PTEDGSEEPG
SETSDAKSTP TAEDVTAPLV DEGAPGKQAA AQPHTEIPEG TTAEEAGIGD TPSLEDEAAG
HVTQARMVSK SKDGTGSDDK KAKGADGKTK IATPRGAAPP GQKGQANATR IPAKTPPAPK
TPPSSGEPPK SGDRSGYSSP GSPGTPGSRS RTPSLPTPPT REPKKVAVVR TPPKSPSSAK
SRLQTAPVPM PDLKNVKSKI GSTENLKHQP GGGKVQIINK KLDLSNVQSK CGSKDNIKHV
PGGGSVQIVY KPVDLSKVTS KCGSLGNIHH KPGGGQVEVK SEKLDFKDRV QSKIGSLDNI
THVPGGGNKK IETHKLTFRE NAKAKTDHGA EIVYKSPVVS GDTSPRHLSN VSSTGSIDMV
DSPQLATLAD EVSASLAKQG L

FIB1
MGRPEFNRGG GGGGFRGGRG GDRGGSRGGF GGGGRGGYGG GDRGSFGGGD RGGFRGGRGG
GDRGGFRGGR GGGDRGGFGG RGSPRGGFGG RGSPRGGRGS

Edc3
PSTNSTKLKS AETYSSKNKW SMDCDEEFDF AANLEKFDKK QVFAEFREKD KKDPAKLLVS
HNKSPNRNYH HKQNVLGPSV KDEFVDLPSA GSQINGIDAV LSSSSNGHVT PGSKKGSRET
LKK

Ddx4
MGDEDWEAEI NPHMSSYVPI FEKDRYSGEN GDNFNRTPAS SSEMDDGPSR RDHFMKSGFA
SGRNFGNRDA GECNKRDNTS TMGGFGVGKS FGNRGFSNSR FEDGDSSGFW RESSNDCEDN
PTRNRGFSKR GGYRDGNNSE ASGPYRRGGR GSFRGCRGGF GLGSPNNDLD PDECMQRTGG
LFGSRRPVLS GTGNGDTSQS RSGSGSERGG YKGLNEEVIT GSGKNSWKSE AEGGES

RNase CTD
TGVLEGTTHV CEHCEGTGRV RSVESSALAA LRAVEAEALK GSGSVILKVS RSVGLYILNE
KRDYLQRLLT THGLFVSVVV DDSLHAGDQE IERTELGERI AVAPPPFVEE DDDFDPNAYD
DEEEEDDVIL DDEDDTDRED TDDDDATTRK SARDDERGDR KGRRGRRDRN RGRGRRDERD
GETESEDEDV VAEGADEDRG EFGDDDEGGR RRRRRGRRGG RRGGREDGDR PTDAFVWIRP
RVPFGENVFT WHDPAALVGG GESRRQAPEP RVDAATEAAP RPERAEREER PGRERGRRGR
DRGRRQRDEA PVAEMTSVES ATVEAAEPFE APILAPPVIA GPPADVWVEL PEVEEAPKKP
KRSRARGKKA TETSVEAIDT VTEVAAEAPA PETAEPEAVE VAPPAPTVEA APEPGPVVEA
VEEAQPAEPD PNEITAPPEK PRRGWWRR

1.2. **Sequence patterning descriptors.** Here we briefly show the equations for the two variants of sequence charge decoration $SCD$ and refer the readers to the original literature for details.[1, 2] The variant of $SCD$ in the low salt regime is defined as

$$SCD_{\text{lowsalt}} = N^{-1}\Sigma_i\Sigma_{j,j>i}q_iq_j(j-i), \tag{S1}$$

whereas the variant at high salt regime is defined as

$$SCD_{\text{highsalt}} = N^{-1}\Sigma_i\Sigma_{j,j>i}q_iq_j(j-i)^{1/2}, \tag{S2}$$

in which $q_i$ and $q_j$ are the charges of the $i$-th and $j$-th amino acids in the sequence.

As an extension to the sequence hydropathy decoration $SHD$,[3] we can define a salt-dependent variant of $SHD$ as

$$SHD_{\text{salt}}(C) = N^{-1}\Sigma_i\Sigma_{j,j>i}(\lambda_i'(C)+\lambda_j'(C))|j-i|^{\beta}, \tag{S3}$$

in which $\lambda_i'(C)$ and $\lambda_j'(C))$ are the salt-dependent hydropathy scales of the $i$-th and $j$-th amino acids defined in Eq. 2 of the main text.

1.3. **HPS model.** There are three types of interactions, including bonded, electrostatic and short-range pairwise interaction terms characterized by amino acid hydropathy.[4] Bonded interactions are modeled by a harmonic potential with a spring constant of 10 kJ/Å$^2$ and a bond length of 3.8 Å. Electrostatic interactions are modeled using a Coulombic term with Debye-Hückel [5] electrostatic screening,

$$E_{ij}(r) = \frac{q_i q_j}{4\pi D r} \exp(-r/\kappa), \tag{S4}$$

in which $\kappa$ is the Debye screening length and $D = 80$, the dielectric constant of the solvent. The short-range pairwise interaction is modeled using Ashbaugh-Hatch functional form[6],

$$\Phi(r) = \begin{cases} \Phi_{LJ} + (1-\lambda)\epsilon, & \text{if } r \leq 2^{1/6}\sigma \\ \lambda \Phi_{LJ}, & \text{otherwise} \end{cases} \tag{S5}$$

in which $\Phi_{LJ}$ is the standard Lennard-Jones potential

$$\Phi_{LJ} = 4\epsilon \Big[ \Big(\frac{\sigma}{r}\Big)^{12} - \Big(\frac{\sigma}{r}\Big)^{6} \Big]. \tag{S6}$$

The $\lambda$ value in the pairwise interaction term is the arithmetic average of the $\lambda$ values of the two amino acids. The amino-acid specific parameters of the model are shown in Table S2. The interaction strength $\epsilon$ is set to 0.16 kcal/mol for E-cad simulations following the recent work to best match the experimental FRET data,[7] and 0.2 kcal/mol for all the other simulations since this was the optimal value for a variety of IDP sequences.[8]

1.4. **Molecular dynamics simulations.** The HOOMD-Blue software v2.9.3 [9] together with the azplugins [10] were used for running the molecular dynamics simulations. All simulations were run using a Langevin thermostat with a friction coefficient of 0.01 ps$^{-1}$, a time step of 10 fs and a temperature of 298 K. The simulations of the 17 sequences with experimental LLPS behaviors were run for 5 $\mu$s and the first 0.5 $\mu$s was dumped as equilibration. The simulations of the 530 sequences from the Disprot database were run for 0.5 $\mu$s and the first 0.1 $\mu$s was dumped as equilibration. For both LLPS-enabled and Disprot sequences, simulations were run at six different salt concentrations including 0.01, 0.05, 0.1, 0.5, 1 and 2 M. All the simulations for parameterizing salting-out term using E-cad data were run for 5 $\mu$s and the first 0.5 $\mu$s was dumped as equilibration at salt concentrations 0.022, 0.062, 0.112, 0.212, 0.512, 1.012 and 2.012 M to be consistent with the experimental conditions.[7] We note these simulations are sufficient long to simulate $R_\text{g}$ with less than 1% errors, we therefore do not include the errorbars in all the $R_\text{g}$ plots.

#### 2. Supplementary Figures

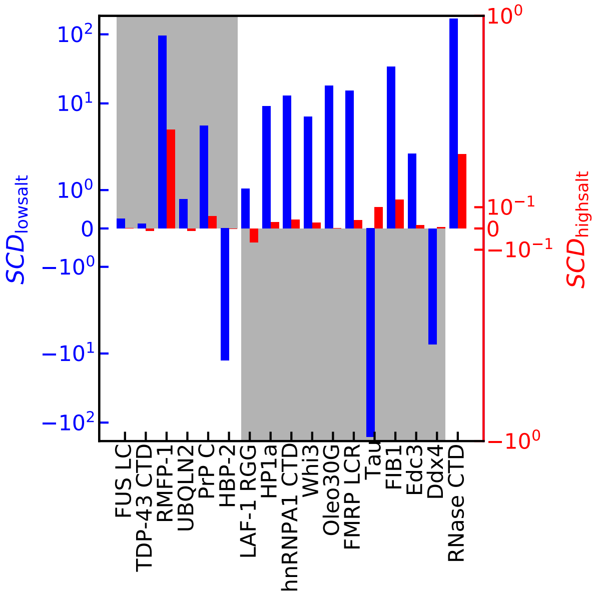


Figure S1. $SCD_{\mathrm{lowsalt}}$ (blue) and $SCD_{\mathrm{highsalt}}$ (red) for the 17 sequences with salt-dependent LLPS behaviors. The gray shading shows the expected behaviors on these values using experimentally observed salt-dependent phase behaviors.

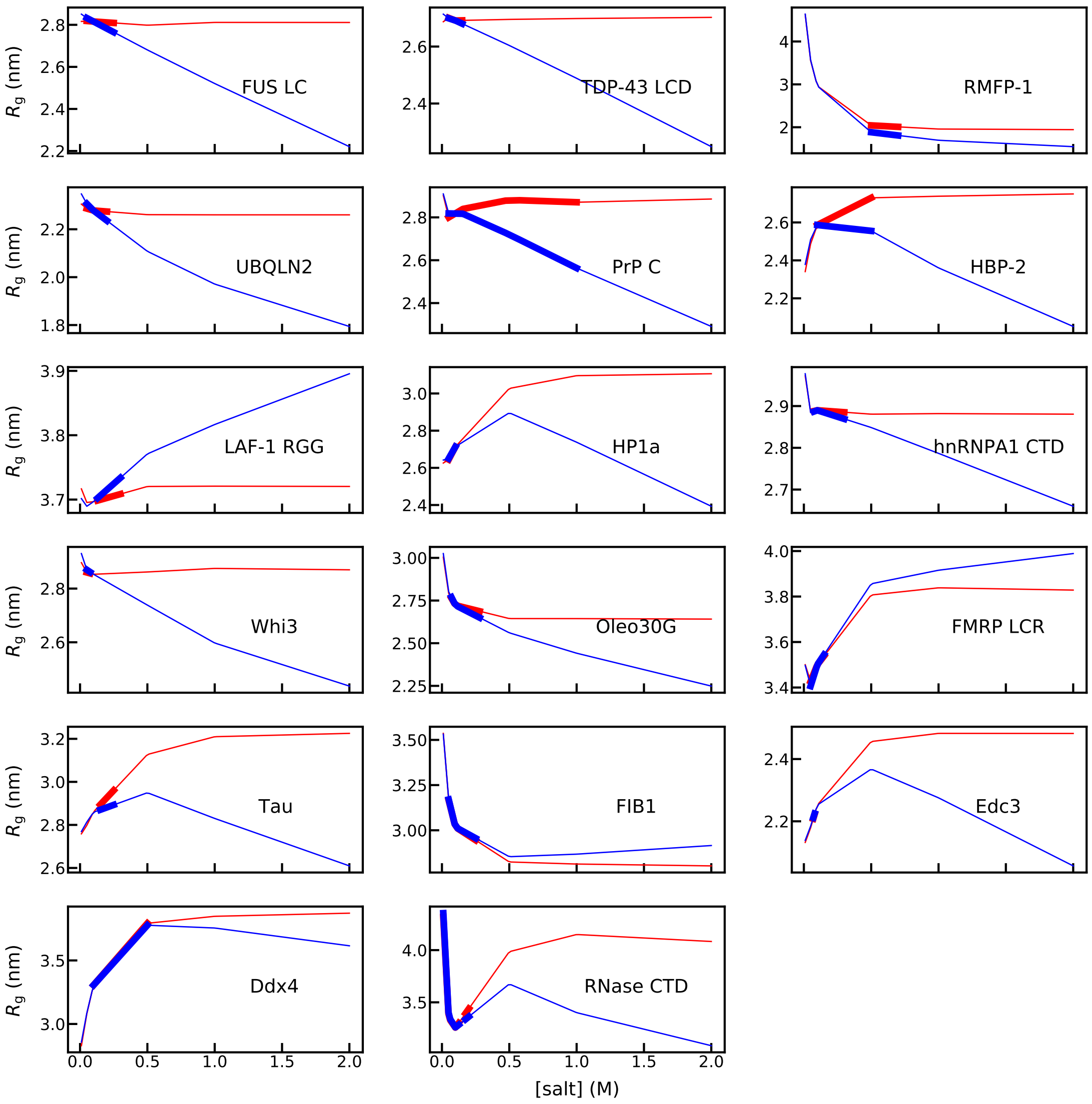


FIGURE S2. $R_g$ from the HPS-salt model (blue) and the HPS model (red) for the 17 sequences using an interaction strength of $\epsilon$=0.2 kcal/mol. The experimental ranges of salt concentrations are highlighted in thick lines.

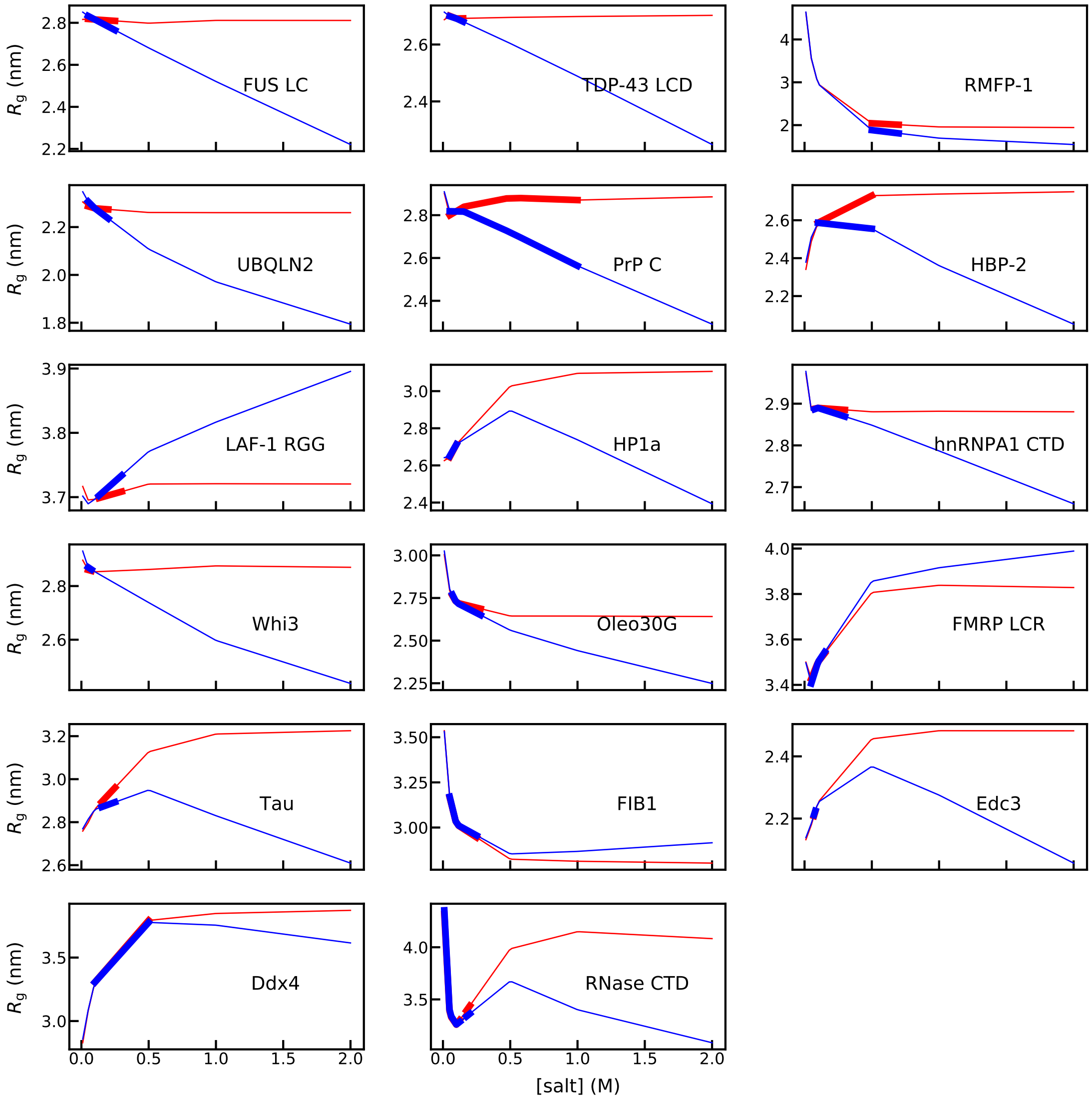


Figure S3. $R_g$ from the HPS-salt model (blue) and the HPS model (red) for the 17 sequences using an interaction strength of $\epsilon$=0.16 kcal/mol. The experimental ranges of salt concentrations are highlighted in thick lines.

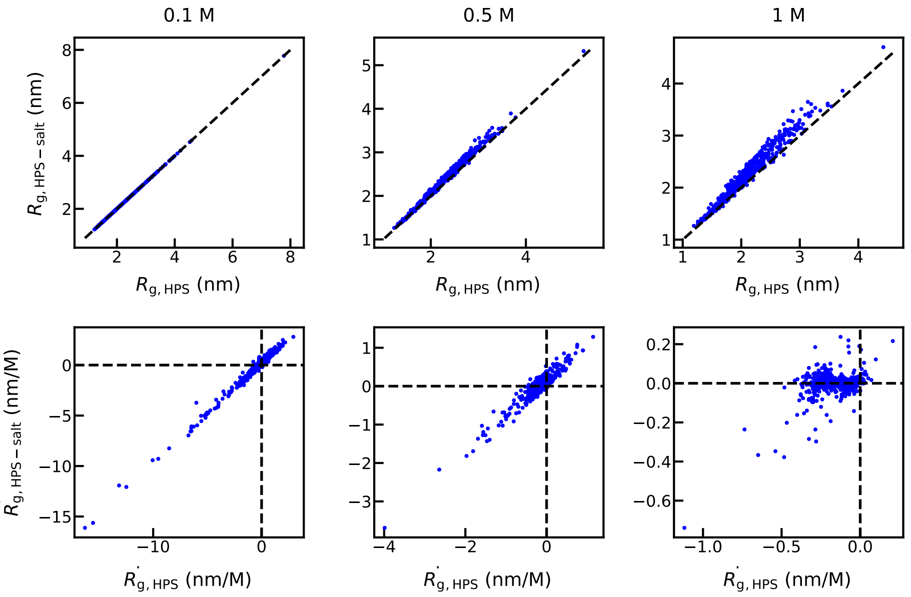


Figure S4. Top: Comparison between $R_g$ from HPS-salt and HPS models at different salt concentrations for the IDP sequences from the Disprot database.[11, 12] Bottom: Comparision between the derivative of $R_g$ along the salt concentration ($\dot{R}_g$) from the HPS-salt and HPS models.

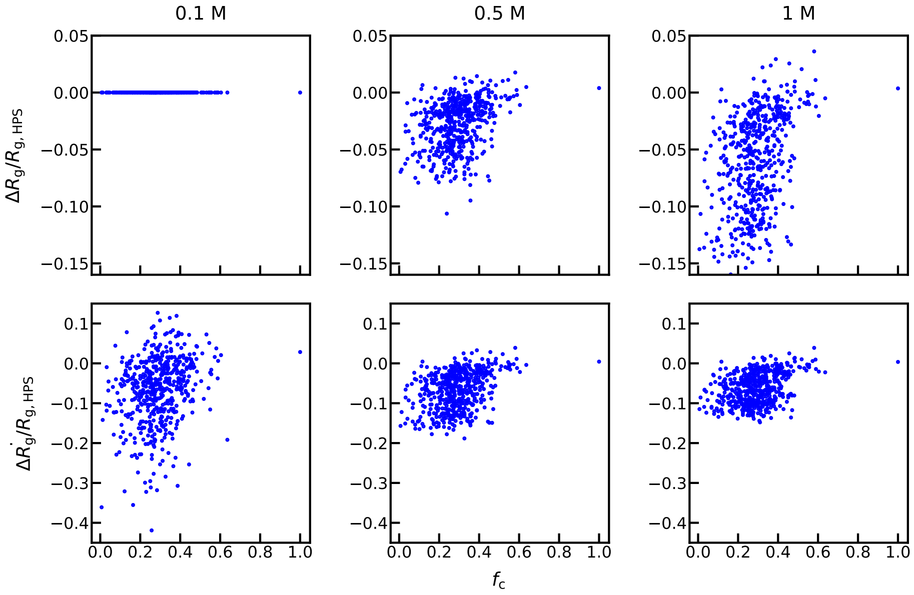


FIGURE S5. Top: Difference between $R_g$ from HPS-salt and HPS models at different salt concentrations for the IDP sequences from the Disprot database.[11, 12] Bottom: Difference between the derivative of $R_g$ along the salt concentration ($\dot{R}_g$) from the HPS-salt and HPS models. All values are normalized by the $R_g$ from the HPS model at the corresponding salt concentration.

#### 3. Supplementary Tables

Table S1. List of proteins with experimentally observed salt-dependent phase behaviors. We consider close to zero $SCD_{\text{lowsalt}}$ (<0.3) and $\dot{R}_g$ (<0.1 nm/M) as the cases without an obvious salt-dependent behavior (→). * notes the sequence with the disordered region predicted using PONDR.[13]

| Name | Expt | salt range (mM) | $SCD_{\text{lowsalt}}$ | $SHD$ | HPS | HPS-salt |
|---|---|---|---|---|---|---|
| FUS LC | HSPS [14] | 50-250 | → ✗ | ↑ ✓ | → ✗ | ↑ ✓ |
| TDP-43 CTD | HSPS [15] | 50-150 | → ✗ | ↑ ✓ | → ✗ | ↑ ✓ |
| RMFP-1 | HSPS [16] | 500-700 | ↑ ✓ | ↑ ✓ | ↑ ✓ | ↑ ✓ |
| UBQLN2 | HSPS [17] | 150-250 | ↑ ✓ | ↑ ✓ | → ✗ | ↑ ✓ |
| PrP C | HSPS [18] | 31.5-1000 | ↑ ✓ | ↑ ✓ | → ✗ | ↑ ✓ |
| HBP-2 | HSPS [19] | 100-500 | ↓ ✗ | ↑ ✓ | ↓ ✗ | ↑ ✓ |
| HSPS correctness | | | 3/6 | 6/6 | 1/6 | 6/6 |
| LAF-1 RGG | LSPS [20] | 130-300 | ↑ ✗ | ↓ ✓ | → ✗ | ↓ ✓ |
| HP1a | LSPS [21] | 25-100 | ↑ ✗ | ↑ ✗ | ↓ ✓ | ↓ ✓ |
| hnRNPA1 CTD | LSPS [22] | 75-300 | ↑ ✗ | ↑ ✗ | → ✗ | ↑ ✗ |
| Whi3 (231-600*) | LSPS [23] | 20-75 | ↑ ✗ | ↑ ✗ | ↑ ✗ | ↑ ✗ |
| Oleo30G | LSPS [24] | 70-280 | ↑ ✗ | ↑ ✗ | ↑ ✗ | ↑ ✗ |
| FMRP LCR | LSPS [25] | 50-150 | ↑ ✗ | ↓ ✓ | ↓ ✓ | ↓ ✓ |
| Tau | LSPS [26] | 150-250 | ↓ ✓ | ↑ ✗ | ↓ ✓ | ↓ ✓ |
| FIB1 | LSPS [27] | 50-250 | ↑ ✗ | ↓ ✓ | ↑ ✗ | ↑ ✗ |
| Edc3 | LSPS [28] | 70-80 | ↑ ✗ | ↑ ✗ | ↓ ✓ | ↓ ✓ |
| Ddx4 | LSPS [29] | 100-500 | ↓ ✓ | ↑ ✗ | ↓ ✓ | ↓ ✓ |
| LSPS correctness | | | 2/10 | 3/10 | 5/10 | 6/10 |
| RNase CTD | MSPS [30] | 0-125,175-200 | ↑ ✗ | ↑ ✗ | ↑+↓ ✓ | ↑+↓ ✓ |
| MSPS correctness | | | 0/1 | 0/1 | 1/1 | 1/1 |
| Total correctness | | | 5/17 | 9/17 | 7/17 | 13/17 |

TABLE S2. The amino acid parameters used in the model. $\sigma$ is the diameter of the amino acid used in the short-ranged pair potential. $\lambda$ is the scaled hydropathy from the literature [4]. $k_s$ is the salting-out constants from literature [31, 32, 33, 34] and from its correlation with the hydropathy scale (Fig. 2A).

| Type | Mass (amu) | Charge | $\sigma$ (Å) | $\lambda$ | $k_s$ ($M^{-1}$) |
|---|---|---|---|---|---|
| ALA | 71.08 | 0 | 5.04 | 0.730 | -0.010 |
| ARG | 156.20 | 1 | 6.56 | 0.000 | -0.255 |
| ASN | 114.10 | 0 | 5.68 | 0.432 | -0.155 |
| ASP | 115.10 | -1 | 5.58 | 0.378 | -0.090 |
| CYS | 103.10 | 0 | 5.48 | 0.595 | -0.140 |
| GLN | 128.10 | 0 | 6.02 | 0.514 | -0.095 |
| GLU | 129.10 | -1 | 5.92 | 0.459 | -0.070 |
| GLY | 57.05 | 0 | 4.50 | 0.649 | -0.040 |
| HIS | 137.10 | 0.5 | 6.08 | 0.514 | -0.081 |
| ILE | 113.20 | 0 | 6.18 | 0.973 | 0.080 |
| LEU | 113.20 | 0 | 6.18 | 0.973 | 0.080 |
| LYS | 128.20 | 1 | 6.36 | 0.514 | -0.081 |
| MET | 131.20 | 0 | 6.18 | 0.838 | 0.030 |
| PHE | 147.20 | 0 | 6.36 | 1.000 | 0.070 |
| PRO | 97.12 | 0 | 5.56 | 1.000 | 0.085 |
| SER | 87.08 | 0 | 5.18 | 0.595 | -0.053 |
| THR | 101.10 | 0 | 5.62 | 0.676 | -0.025 |
| TRP | 186.20 | 0 | 6.78 | 0.946 | 0.070 |
| TYR | 163.20 | 0 | 6.46 | 0.865 | 0.070 |
| VAL | 99.07 | 0 | 5.86 | 0.892 | 0.060 |

#### References


[1] Sawle, L.; Ghosh, K. *J. Chem. Phys.* **2015**, *143*, 085101.

[2] Huihui, J.; Firman, T.; Ghosh, K. *J. Chem. Phys.* **2018**, *149*, 085101.

[3] Zheng, W.; Dignon, G.; Brown, M.; Kim, Y. C.; Mittal, J. *J. Phys. Chem. Lett.* **2020**, *11*, 3408–3415.

[4] Kapcha, L. H.; Rossky, P. J. *J. Mol. Biol.* **2014**, *426*, 484–498.

[5] Debye, P.; Hückel, E. *Physikalische Zeitschrift* **1923**, *24*, 185–206.

[6] Ashbaugh, H. S.; Hatch, H. W. *J. Am. Chem. Soc.* **2008**, *130*, 9536–9542.

[7] Wiggers, F.; Wohl, S.; Dubovetskyi, A.; Rosenblum, G.; Zheng, W.; Hofmann, H. *bioRxiv* **2021**, https://www.biorxiv.org/content/10.1101/2021.02.03.429507v1.

[8] Dignon, G. L.; Zheng, W.; Kim, Y. C.; Best, R. B.; Mittal, J. *PLoS Comput. Biol.* **2018**, *14*, e1005941.

[9] Anderson, J. A.; Glaser, J.; Glotzer, S. C. *Comput. Mater. Sci.* **2020**, *173*, 109363.

[10] https://github.com/mphowardlab/azplugins.

[11] Sickmeier, M.; Hamilton, J. A.; LeGall, T.; Vacic, V.; Cortese, M. S.; Tantos, A.; Szabo, B.; Tompa, P.; Chen, J.; Uversky, V. N.; Obradovic, Z.; Dunker, A. K. *Nucleic Acids Res.* **2006**, *35*, D786–D793.

[12] Piovesan, D.; Tabaro, F.; Mičetić, I.; Necci, M.; Quaglia, F.; Oldfield, C. J.; Aspromonte, M. C.; Davey, N. E.; Davidović, R.; Dosztányi, Z.; et al. *Nucleic Acids Res.* **2017**, *45*, D219–D227.

[13] Li, X.; Romero, P.; Rani, M.; Dunker, A. K.; Obradovic, Z. *Genome Informatics* **1999**, *10*, 30–40.

[14] Burke, K. A.; Janke, A. M.; Rhine, C. L.; Fawzi, N. L. *Mol. Cell* **2015**, *60*, 231–241.

[15] Conicella, A. E.; Zerze, G. H.; Mittal, J.; Fawzi, N. L. *Structure* **2016**, *24*, 1537–1549.

[16] Kim, S.; Yoo, H. Y.; Huang, J.; Lee, Y.; Park, S.; Park, Y.; Jin, S.; Jung, Y. M.; Zeng, H.; Hwang, D. S.; et al. *ACS Nano* **2017**, *11*, 6764–6772.

[17] Yang, Y.; Jones, H. B.; Dao, T. P.; Castañeda, C. A. *J. Phys. Chem. B* **2019**, *123*, 3618–3629.

[18] Kostylev, M. A.; Tuttle, M. D.; Lee, S.; Klein, L. E.; Takahashi, H.; Cox, T. O.; Gunther, E. C.; Zilm, K. W.; Strittmatter, S. M. *Mol. Cell* **2018**, *72*, 426–443.

[19] Le Ferrand, H.; Duchamp, M.; Gabryelczyk, B.; Cai, H.; Miserez, A. *J. Am. Chem. Soc.* **2019**, *141*, 7202–7210.

[20] Elbaum-Garfinkle, S.; Kim, Y.; Szczepaniak, K.; Chen, C. C.-H.; Eckmann, C. R.; Myong, S.; Brangwynne, C. P. *Proc. Natl. Acad. Sci. U.S.A.* **2015**, *112*, 7189–7194.

[21] Strom, A. R.; Emelyanov, A. V.; Mir, M.; Fyodorov, D. V.; Darzacq, X.; Karpen, G. H. *Nature* **2017**, *547*, 241–245.

[22] Martin, E. W.; Thomasen, F. E.; Milkovic, N. M.; Cuneo, M. J.; Grace, C. R.; Nourse, A.; Lindorff-Larsen, K.; Mittag, T. *Nucleic Acids Res.* **2021**, *49*, 2931–2945.

[23] Zhang, H.; Elbaum-Garfinkle, S.; Langdon, E. M.; Taylor, N.; Occhipinti, P.; Bridges, A. A.; Brangwynne, C. P.; Gladfelter, A. S. *Mol. Cell* **2015**, *60*, 220–230.

[24] Reed, E. H.; Hammer, D. A. *Soft Matter* **2018**, *14*, 6506–6513.

[25] Tsang, B.; Arsenault, J.; Vernon, R. M.; Lin, H.; Sonenberg, N.; Wang, L.-Y.; Bah, A.; Forman-Kay, J. D. *Proc. Natl. Acad. Sci. U.S.A.* **2019**, *116*, 4218–4227.

[26] Hernández-Vega, A.; Braun, M.; Scharrel, L.; Jahnel, M.; Wegmann, S.; Hyman, B. T.; Alberti, S.; Diez, S.; Hyman, A. A. *Cell Rep.* **2017**, *20*, 2304–2312.

[27] Feric, M.; Vaidya, N.; Harmon, T. S.; Mitrea, D. M.; Zhu, L.; Richardson, T. M.; Kriwacki, R. W.; Pappu, R. V.; Brangwynne, C. P. *Cell* **2016**, *165*, 1686–1697.

[28] Schütz, S.; Nöldeke, E. R.; Sprangers, R. *Nucleic Acids Res.* **2017**, *45*, 6911–6922.

[29] Brady, J. P.; Farber, P. J.; Sekhar, A.; Lin, Y.-H.; Huang, R.; Bah, A.; Nott, T. J.; Chan, H. S.; Baldwin, A. J.; Forman-Kay, J. D.; et al. *Proc. Natl. Acad. Sci. U.S.A.* **2017**, *114*, E8194–E8203.

[30] Al-Husini, N.; Tomares, D. T.; Bitar, O.; Childers, W. S.; Schrader, J. M. *Mol. Cell* **2018**, *71*, 1027–1039.

[31] Schrier, E. E.; Schrier, E. B. *J. Phys. Chem.* **1967**, *71*, 1851–1860.

[32] Nandi, P. K.; Robinson, D. R. *J. Am. Chem. Soc.* **1972**, *94*, 1299–1308.

[33] Nandi, P. K.; Robinson, D. R. *J. Am. Chem. Soc.* **1972**, *94*, 1308–1315.

[34] Baldwin, R. L. *Biophys. J.* **1996**, *71*, 2056–2063.
